# Exposure to the predator odor TMT induces early and late differential gene expression related to stress and excitatory synaptic function throughout the brain in male rats

**DOI:** 10.1101/2020.05.26.116657

**Authors:** Ryan E. Tyler, Ben Weinberg, Dennis Lovelock, Laura Ornelas, Joyce Besheer

## Abstract

Persistent changes in brain stress and glutamatergic function are associated with post-traumatic stress disorder (PTSD). Rodent exposure to the predator odor trimethylthiazoline (TMT) is an innate stressor that produces lasting behavioral consequences relevant to PTSD. As such, the goal of the present study was to assess early (6 hours and 2 days) and late (4 weeks) changes to gene expression (RT-PCR) related to stress and excitatory function following TMT exposure in male, Long-Evans rats. During TMT exposure, rats engaged in stress reactive behaviors, including digging and immobility. Further, the TMT group displayed enhanced exploration and mobility in the TMT-paired context one week after exposure, suggesting a lasting contextual reactivity. Gene expression analyses revealed upregulated *FKBP5* 6 hours post-TMT in the hypothalamus and dorsal hippocampus. Two days after TMT, *GRM3* was downregulated in the prelimbic cortex and dorsal hippocampus, but upregulated in the nucleus accumbens. This may reflect an early stress response (*FKBP5*) that resulted in later glutamatergic adaptation (*GRM3*). Finally, four weeks after TMT exposure, several differentially expressed genes known to mediate excitatory tripartite synaptic function were observed. Specifically in the prelimbic cortex (*GRM5, DLG4* and *SLC1A3* upregulated), infralimbic cortex (*GRM2* downregulated, *Homer1* upregulated), nucleus accumbens (*GRM7* and *SLC1A3* downregulated), dorsal hippocampus (*FKBP5* and *NR3C2* upregulated, *SHANK3* downregulated) and ventral hippocampus (*CNR1, GRM7, GRM5, SHANK3*, and *Homer1* downregulated). These data demonstrate that TMT exposure stress induces early and late stress and excitatory molecular adaptations, which may help us understand the persistent glutamatergic dysfunction observed in PTSD.

## Introduction

Post-traumatic stress disorder (PTSD) is a neuropsychiatric disorder that develops in some individuals after experiencing or witnessing trauma^1^. The American Psychiatric Association Diagnostic and Statistical Manual of Mental Disorders (DSM-5) characterizes PTSD within four symptom clusters including re-experiencing (flashbacks, unwanted memories), avoidance (external reminders), hyperarousal/hypervigilance, and negative mood/thoughts^2^. In 2013, the lifetime prevalence of PTSD was reported at 6.1%, which was a total of 14.4 million Americans^3^. Unfortunately, PTSD is a debilitating and enduring disorder with limited treatment options^3^. Therefore, there exists an urgent need to better understand the biological mechanisms underlying PTSD.

Clinical studies show dysregulated HPA-axis function in patients with PTSD^4, 5^. For example, some studies report lower baseline cortisol levels in individuals with PTSD compared to trauma-exposed controls without PTSD or controls not reporting a traumatic experience or PTSD^6,7,8,9^. Additionally, individuals with PTSD show changes in glucocorticoid receptor (GR) expression, binding to glucocorticoids, and associated GR polymorphisms^10, 11, 12^. Binding of cortisol to the GR downregulates corticotropin releasing factor (CRF) expression in the hypothalamus, acting as a negative feedback of HPA-axis activation^13^. Interestingly, in PTSD participants, the FK506 binding protein 5 (FKBP5), a chaperone protein that affects GR and mineralocorticoid receptor (MR) binding sensitivity for glucocorticoids and inhibits cellular translocation, is differentially expressed in several RNA sequencing studies compared to control participants both in peripheral blood cells, and post-mortem PFC brain tissue^14, 15, 16^. Together, these data provide strong support for the hypothesis that PTSD is associated with a persistent dysregulation in the canonical HPA-axis stress system^4^.

In addition to HPA-axis dysfunction, multiple lines of evidence suggest a role for dysregulated excitatory signaling in PTSD^17^. For example, studies have reported changes in brain and blood glutamate concentrations in individuals with PTSD compared to control participants^18, 19^. A positron emission tomography (PET) study showed that individuals with PTSD exhibit enhanced metabotropic glutamate receptor 5 (mGluR5) availability throughout the cortex^20, 21^. Furthermore, this increased mGluR5 availability positively correlated with avoidance symptom severity in the PTSD group^20^. Another PET study revealed increased endocannabinoid receptor type 1 (CB1) availability associated with PTSD^22^, which is interesting because CB1 receptors are inhibitory presynaptic receptors that are functionally linked to post-synaptic mGluR5 signaling^23^. Finally, preclinical studies show that stress-induced glucocorticoid release modulates multiple aspects of glutamatergic synaptic transmission^24^, suggesting the possibility that HPA-axis and excitatory dysfunctions associated with PTSD are related.

Animal models of traumatic stress and PTSD have used exposure to the scent of a predator as a stressor that produces lasting behavioral consequences with relevance to traumatic disorders^25, 26, 27, 28,29,30,31,32,33^. For example, exposure to the synthetically derived fox odor 2,5-dihydro-2,4,5-trimethylthiazoline (TMT) produced avoidance and freezing behaviors, indicative of a fear-like response, and increased serum corticosterone, indicative of a stress response^26,27^. Other studies have shown lasting anxiety-like and hyperarousal behavior ^31^, as well as avoidance of a TMT-paired context or cue^32^. Additionally, 24 days after a single TMT exposure, re-exposure to the TMT-paired context elicited differential expression of metabotropic glutamate receptor sub-type 5 (*GRM5)* gene expression in the amygdala and mPFC in a behaviorally-defined “resilient” group compared to a “susceptible” and control group^31^. Together, these data show that predator odor exposure produces behavioral and molecular changes indicative of a stress response, and a lasting behavioral profile capable of modeling some aspects of PTSD symptomology.

The goal of the present work was to assess early (6 hours and 2 days) and late (4 weeks) gene expression changes throughout the brain following TMT stress exposure. Gene expression analyses focused on stress/HPA-axis function and excitatory neurotransmission-related targets supported by clinical data^1, 16, 17, 20^. Stress-related genes included *NR3C1* (encodes for GR), *NR3C2* (encodes for MR), *FKBP5*, and *CRF*. Excitatory-related genes included several metabotropic glutamate receptors (*GRM2, GRM3, GRM5, GRM7*), *CNR1* (encodes for CB1), and the *SLC1A3* gene (encodes for EAAT-1). Additionally, changes in *GRM5* were followed up by examining genes related to the excitatory post-synaptic density including *DLG4* (encodes for PSD-95), *Homer1*, and *SHANK3*^34^. Gene expression was examined in the prelimbic cortex, infralimbic cortex, nucleus accumbens, hypothalamus, amygdala, dorsal hippocampus, and ventral hippocampus because analogous brain regions are implicated in PTSD and stress^35, 36^. Time points were chosen to capture both relatively immediate (6 hours, 2 days) and late (4 weeks) effects of TMT on gene expression. Most genes were evaluated in all brain regions at each time point following TMT exposure to provide a comprehensive and dynamic assessment of excitatory- and stress-related gene expression changes. In addition to examination of gene expression, behavior during the TMT exposure, and upon context re-exposure one week after TMT were examined to assess reactivity to TMT exposure and the TMT-paired context. Together, these data could inform our understanding of the molecular and behavioral changes following an innate stressor.

## Methods

### Animals

Male Long-Evans rats (n=52; Envigo, Indianapolis, IN) were used for all experiments. Rats arrived at 7 weeks and were handled for at least one min for one week prior to experiments. Rats were housed in ventilated cages with access to food and water ad libitum and maintained in a temperature and humidity controlled vivarium with a 12 hour light/dark cycle. All experiments were conducted during the light cycle. Rats were under continuous care and monitoring by veterinary staff from the Division of Comparative Medicine at UNC-Chapel Hill. All procedures were conducted in accordance with the NIH Guide to Care and Use of Laboratory Animals and institutional guidelines.

*Experiment 1: Assessment of gene expression 6 hours and 2 days after TMT exposure*

### TMT Exposure

Rats were transported from the vivarium in the home cage to a separate, well-ventilated room that contained the test chambers in which rats were exposed to TMT (45.72 × 17.78 × 21.59 cm; UNC Instrument Shop, Chapel Hill, NC). The length of the back wall of the test chambers was opaque white with two opaque black side walls and a clear, plexiglass front wall to enable video recordings, and a clear sliding lid. A small, metal basket was hung on the right-side wall (17.8 cm above the floor) to hold a piece of filter paper. Rats were placed in the test chamber for 10 min on two consecutive days before the experiment for habituation. On the TMT exposure day, rats were placed in the chambers for 20 mins. During this session, 10 µl of TMT (2,5-dihydro-2,4,5-trimethylthiazoline, 97% purity; SRQ Bio, Sarasota, FL) or water for controls was pipetted onto the filter paper in the metal baskets. The control group was always run before the TMT group to prevent odor contamination. The odor exposure session was video recorded for evaluation of behavior using ANY-maze Video Tracking System (Version 6.12, Stoelting Co. Wood Dale, IL). All animals were transferred from double to single housing after the TMT exposure session. One group of rats was sacrificed 6 hours after the exposure (Control group n=6; TMT group n=12). Another group of rats were sacrificed two days (54 hours) after the exposure (Control group n=6; TMT group n=12). Controls for each time point were pooled for behavioral and gene expression analyses. Groups were staggered such that all rats were sacrificed on the same day.

### Brain tissue collection and sectioning

Animals were anesthetized with sodium pentobarbital (Patterson Veterinary, MA; 100 mg/kg, IP) and perfused with ice cold PBS (0.1 M, Fisher, PA) in order to clear peripheral biomolecules from the brain. After perfusion, brains were rapidly extracted and flash frozen with isopentane (Sigma-Aldrich, MI). Brains were stored at -80°C until brain region sectioning. Brains were sectioned on a cryostat (−20°C) up to a predetermined bregma for each region of interest (ROI) according to^37^. Then, a micropunch tool was used to remove tissue specific to each brain region as illustrated in Table 1. Some ROIs were separated by left and right hemispheres, and all qPCR experiments used the right hemisphere when separated. Brain regions that were not bisected by left and right hemisphere used the entire tissue section for qPCR experiments. A complete list of all tissue excision specifications can be found for all ROIs in Table 1. Brain sections were stored at -80°C until qPCR analysis.

**Table 1.**
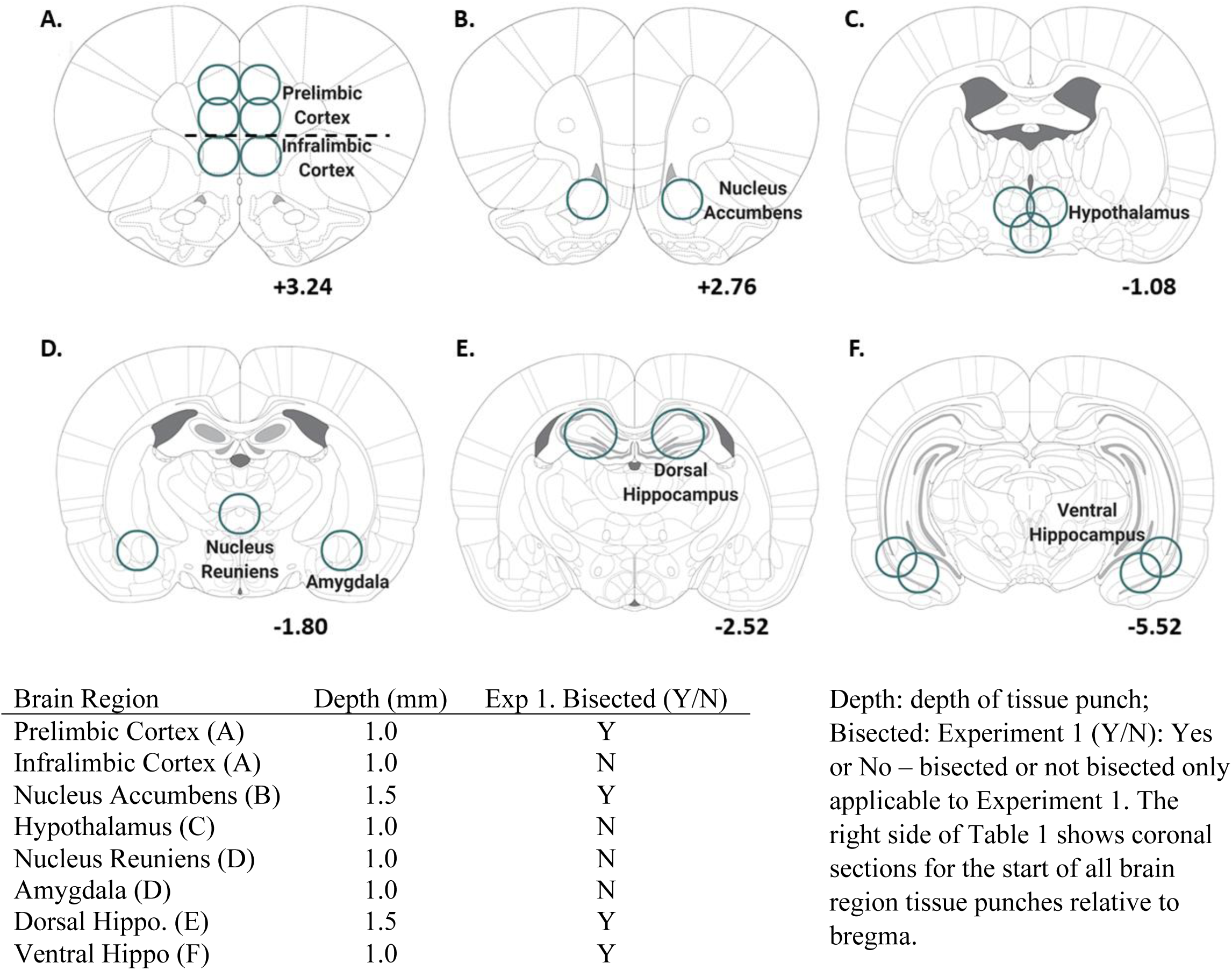
Details of tissue extraction for each brain region.

*Experiment 2: Assessment of gene expression 4 weeks after TMT exposure*

### TMT Exposure

The TMT procedure for this experiment differed slightly from Experiment 1. First, rats were single housed upon arrival, and remained single housed for the duration of the experiment. Second, the TMT exposure was 10 min, rather than 20 min, in duration. Third, white bedding (approximately 600 ml) covered the chamber floor for these experiments to examine digging behavior as an additional behavioral measure. Lastly, animals were not habituated to the chamber environment prior to TMT exposure. Otherwise, the TMT exposure protocol was identical to Experiment 1.

### Context Re-exposure

Seven days after TMT exposure, rats were returned to the chambers in which they had been exposed to TMT (including bedding), but without TMT present. This context re-exposure was 5 min in duration and video recorded for behavioral analyses identical to those assessed during TMT exposure.

### Brain tissue collection

After the conclusion of the context re-exposure test, rats remained undisturbed in their home cage until sacrifice (4 weeks post-TMT exposure; 3 weeks post-context re-exposure). Rats were deeply anesthetized with 5% isoflurane (Baxter Healthcare, NC) in oxygen, followed by rapid brain extraction and flash freezing in isopentane (Sigma-Aldrich, MI). Brains were stored at - 80°C until brain sectioning. ROI location, and micro-punch width and depth were identical to Experiment 1, but both hemispheres were combined for all Experiment 2 RT-PCR analyses. See Table 1 for tissue excision specifications.

### Gene expression analysis using Quantitative Polymerase Chain Reaction (qRT-PCR)

#### Experiments 1 and 2

##### RNA Extraction

RNA was extracted from brain tissue using the RNeasy Mini Kit (Qiagen, Venlo, Netherlands) according to the manufacturer’s instructions. RLT lysis buffer containing β-mercaptoethanol (Sigma Aldrich) was used for tissue homogenization. RNA concentration and purity for each sample were determined using a Spectrophotometer (Nanodrop 2000, ThermoScientific).

##### Reverse Transcription

RNA was reverse transcribed into cDNA using the QuantiNova Reverse Transcription Kit (Qiagen, Venlo, Netherlands) according to the manufacturer’s instructions. Following reverse transcription, all samples were diluted 1:10 with water (200 uL total), and stored at -20°C before RT-PCR experiments.

##### RT-PCR

The StepOnePlus or QuantStudio3 PCR machine (ThermoFisher) was used for all experiments. Importantly, the same machine was used for all experiments conducted on the same brain region and at the same time point. Using a 96-well plate, each sample was run in triplicate using 10 µL total volume per well with the following components: PowerUp Syber green dye (ThermoFisher, containing ROX dye for passive reference), forward and reverse primers (Eton Biosciences Inc., NC, USA), and cDNA template. The PCR was run with an initial activation for 10 mins at 95°C, followed by 40 cycles of the following: denaturation (95°C for 15s), annealing (60°C for 30s), and extension (72°C for 45s). Melt curves were obtained for all experiments to verify synthesis of a single amplicon. All primer sequences are displayed in Table 2.

**Table 2.**
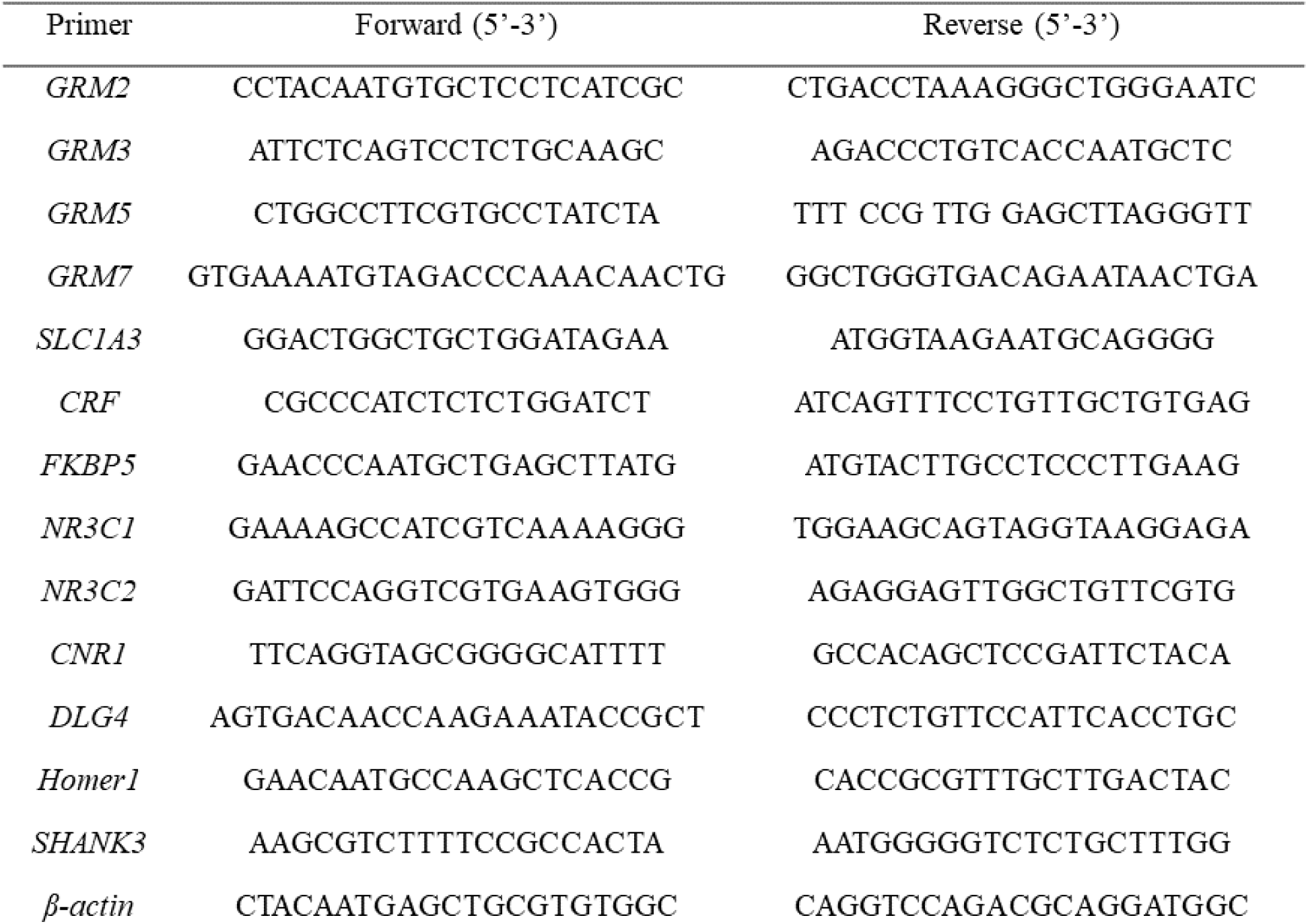
Primer sequences used for RT-PCR experiments.

### Data Analyses

#### Behavioral assessments: TMT exposure and context re-exposure

Using ANY-maze, the length of the rectangular TMT exposure chamber was divided into two compartments for analysis (TMT side and non-TMT side). The basket containing TMT was located on the far end of the TMT side. For Experiment 1, time spent immobile, time spent on the TMT side, distance traveled, and fecal boli during the TMT exposure were assessed. Immobility was operationally defined as the absence of movement other than respiration for longer than 2 seconds. For Experiment 2, time spent digging, time spent immobile, time spent on the TMT side, time spent grooming, distance traveled and fecal boli count were quantified. For Experiment 2, two control and two TMT rats were excluded from the behavioral analyses of TMT exposure due to an error with video recording resulting in 6 control and 6 TMT rats. A two-way RM ANOVA was used to test significant effects of TMT exposure on behavior over time. Sidak’s multiple comparison test was used for all *post hoc* analyses. In Experiment 2, identical dependent measures were analyzed for the context re-exposure test as those analyzed for the TMT exposure. The context re-exposure was not analyzed across time, but as a total of the 5 min test. Supplementary Table 1 displays Experiment 2 behavioral measures (TMT exposure and context re-exposure) in seconds of the total 5 min test. A student’s t-test was used for all two group comparisons. All data are reported as mean ± SEM. Significance was set at p≤0.05.

#### Gene expression assessments

We used the ΔΔCt method to determine fold change relative to controls ^38^. Fold changes were normalized so that average control fold change equaled 1. One-way ANOVAs with *post hoc* Tukey’s multiple comparisons were used for Experiment 1. Experiment 2 data were analyzed using student’s t-tests. Graphs are displayed as transformed fold changes (log2) of controls, from the TMT group only. Zero represents the average fold change of controls. Samples were removed from analysis in case of experimenter error or if determined to be a statistical outlier (greater than 2 std. dev. from the mean). All gene expression data are reported as mean fold change (Supplementary Table 2-4) or the transformed log2 of fold change (Figures) ± SEM. Significance was set at p≤0.05.

## Results

### Experiment 1: Assessment of gene expression 6 hours and 2 days after TMT exposure

#### TMT exposure produces immobility and avoidance behaviors

Figure 1 shows the timeline for Experiment 1 (A), and the behavioral response during TMT exposure (B-D). Analysis of the percent time spent immobile across the 20 min session showed a significant main effect of TMT (F(1, 34)=199.6, p < 0.0001), a significant main effect of time (F(3, 102)=63.7, p < 0.0001), and a significant TMT X time interaction (F(3, 102)=24.5, p<0.0001; Fig. 1B). *Post hoc* analyses showed increased time spent immobile at minute 10, 15, and 20 in the TMT group relative to controls (p<0.05). Total time spent immobile during the 20 min session was 127.9 ± 15.8 s in controls, and 786.2 ± 27.9 s in the TMT group. Analysis of percent time spent on the TMT side showed a main effect of TMT (F(1, 34)=42.8, p < 0.001; Fig. 1C), with the TMT group spending less time on the TMT side. There was no main effect of time or interaction. Total time spent on the TMT side during the 20 min test was 808.0 ± 55.9 s in controls and 180.9 ± 51.1 s in the TMT group. Analysis of distance traveled during exposure yielded a significant main effect of TMT (F (1, 34)=46.3, p < 0.001, Fig. 1D), a significant main effect of time (F(3, 102)=67.1, p < 0.0001), and a significant TMT X time interaction (F(3, 102)=8.3, p < 0.0001). *Post hoc* analyses showed decreased distance traveled in the TMT group compared to the control group at all time points (p<0.05) except the first 5 min. Fecal boli production during the TMT exposure did not differ between the control and TMT groups (Control: 5.9 ± 1.4; TMT: 7.6 ± 0.5; not shown). These data show that TMT exposure elicits a behavioral reactivity characteristic of a fear-like behavior in rats.

**Figure 1.**
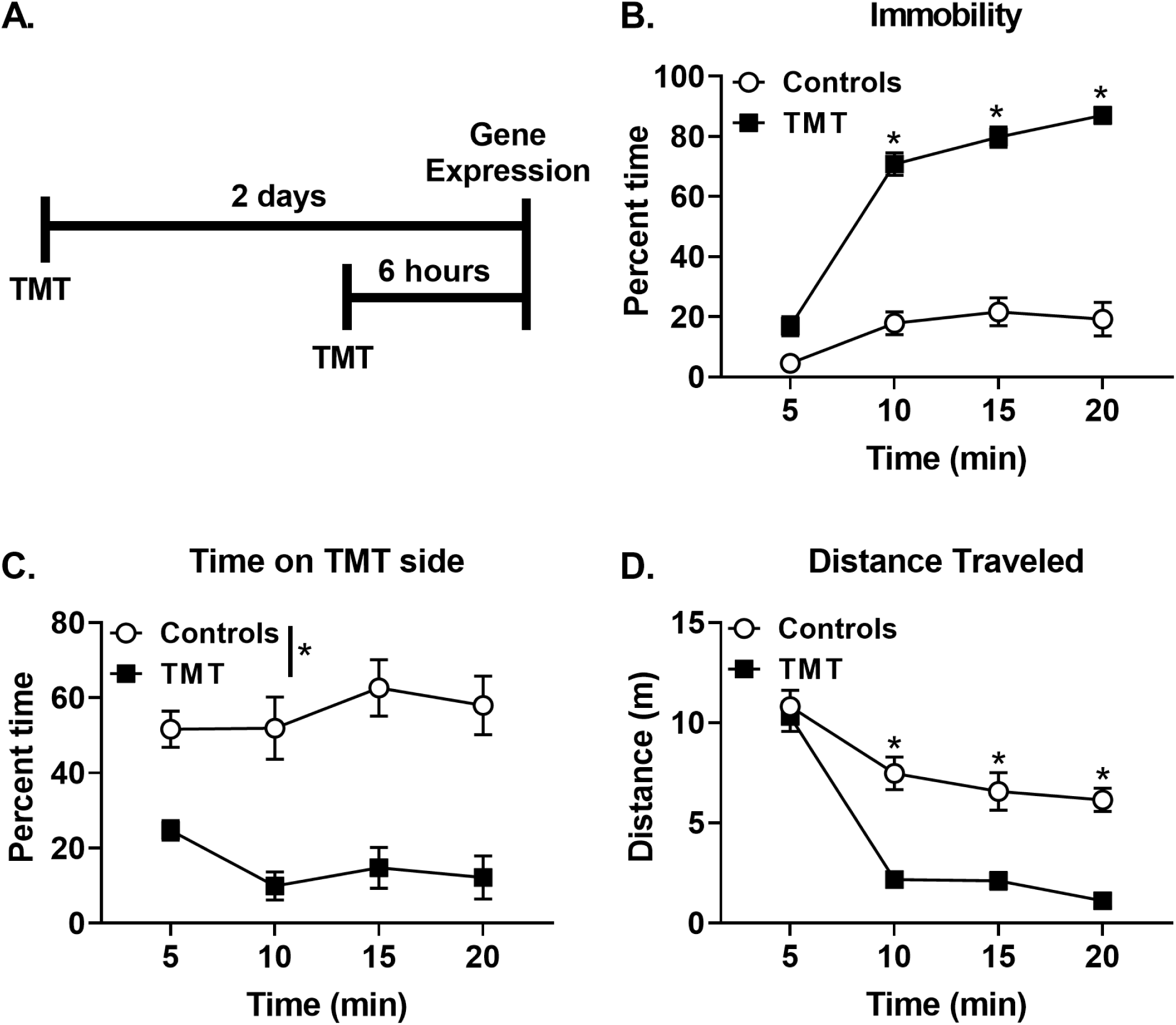
TMT exposure produces freezing and avoidance during exposure. (A) Experimental timeline for Experiment 1. During TMT exposure, the TMT group displayed (B) increased percent time immobile, (C) decreased percent time spent on the TMT side, and (D) reduced distance traveled relative to the Control group. n=12 Control; n=24 TMT. * p≤0.05 significantly different from Control.

#### TMT exposure affects FKBP5 and GRM3 gene expression at early time points

Figure 2 shows gene expression changes 6 hours and 2 days after TMT exposure. In the hypothalamus, *FKBP5* was upregulated (F(2, 33)=3.3, p=0.05, Fig. 2A) six hours after exposure (p<0.05), but not 2 days post-exposure. *FKBP5* expression in the dorsal hippocampus followed the same pattern (F(2, 32)=5.2, p=0.01, *post-hoc* p<0.05, Fig. 2A). In contrast, *FKBP5* expression was not changed in the ventral hippocampus (Fig. 2A). Additionally, Figure 2B shows significant changes in *GRM3* gene expression in the prelimbic cortex, dorsal hippocampus, and nucleus accumbens 2 days following TMT exposure. Specifically, *GRM3* was downregulated in the prelimbic cortex (F(2, 30)=5.0, p=0.01) and dorsal hippocampus (F(2, 30)=3.3, p=0.05), but upregulated in the nucleus accumbens (F(2, 31)=3.7, p=0.04) 2 days following TMT exposure (p<0.05). Additionally, we observed an upwards trend for *CRF* gene expression (p=0.06) 2 days post-TMT (Supplementary Table 3). In contrast, no changes in *NR3C1* or *NR3C2* expression were observed in either dorsal hippocampus or hypothalamus (Supplementary Table 3). *GRM2, GRM5*, or *GRM7* gene expression were not altered following TMT exposure in any of the brain regions examined at these time points (Supplementary Table 2, 3).

**Figure 2.**
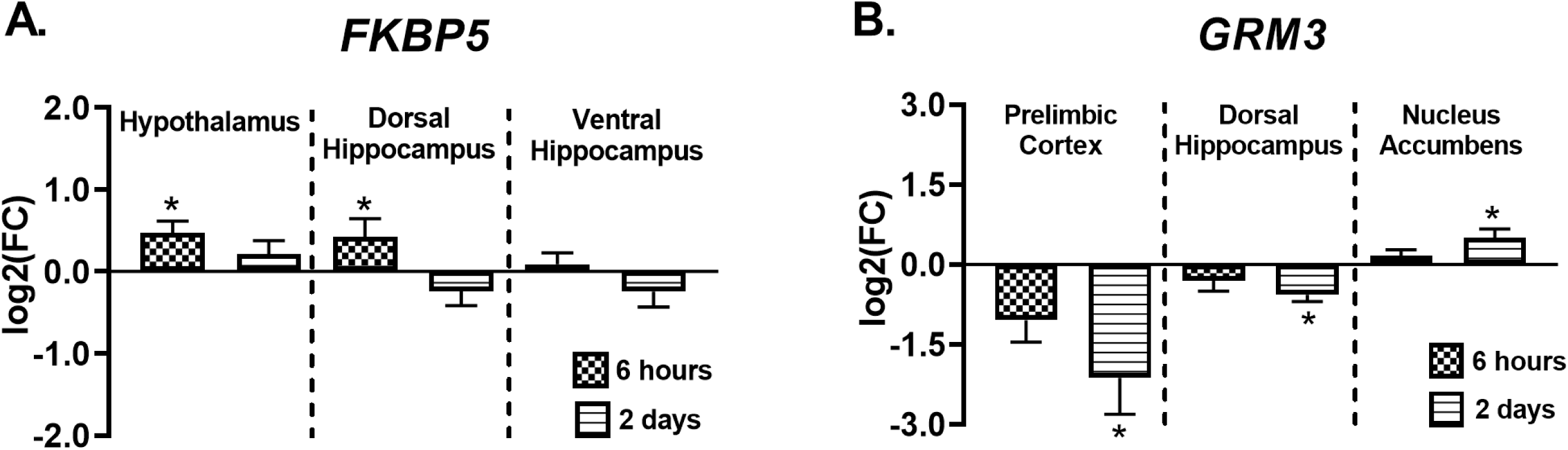
TMT exposure results in differential expression of *FKBP5* and *GRM3*. Six hours after TMT, (A) *FKBP5* gene expression was increased in the hypothalamus (n=12 Control; n=12 TMT) and dorsal hippocampus (n=12 Control; n=11 TMT). Two days after TMT, (B) *GRM3* gene expression was decreased in the prelimbic cortex (n=11 Control; n=12 TMT) and dorsal hippocampus (n=11 Control; n=11 TMT), but upregulated in the nucleus accumbens (n=12 Control; n=12 TMT). * p≤0.05 significantly different from Control.

### Experiment 2: Assessment of gene expression 4 weeks after TMT exposure

#### TMT exposure produces freezing, avoidance, and digging behaviors

Figure 3 shows the experimental timeline (A) for Experiment 2 and the behavioral response during TMT exposure (B-F). Analysis of percent time digging (Fig. 3B) showed a main effect of TMT (F(1,10)=30.2, p=0.0003), a main effect of time (F(4, 40)=36.4, p<0.0001), and a TMT X time interaction (F(4, 40)=33.5, p<0.0001), with increased digging during the first part of the session (min 2, 4, and 6) in the TMT group compared to the control group (p<0.05). For percent time spent immobile (Fig. 3C), there was a significant main effect of TMT (F(1, 10)=208.6, p<0.0001), a significant main effect of time (F(4, 40)=76.1, p<0.0001), and a significant TMT X time interaction (F(4, 40)=76.4, p<0.0001), with increased immobility during the latter part of the session (min 8 and 10) compared to controls (p<0.05). Analysis of percent time spent on the TMT side showed a significant main effect of TMT (F(1, 10)=10.4, p=0.009, Fig. 3D), with the TMT group spending less time on the TMT side compared to controls, but no main effect of time. Analysis of percent time grooming showed a significant main effect of TMT (F(1, 10)=23.5, p=0.0007, Fig. 3E), but no effect of time or interaction. Overall, TMT-exposed rats groomed less than controls. TMT did not significantly affect overall distance traveled (Fig. 3F). Finally, fecal boli production was increased in the TMT group compared to controls (Control: 0.9 ± 0.5; TMT: 5.4 ± 0.8; t(14)=4.8, p=0.0003).

**Figure 3.**
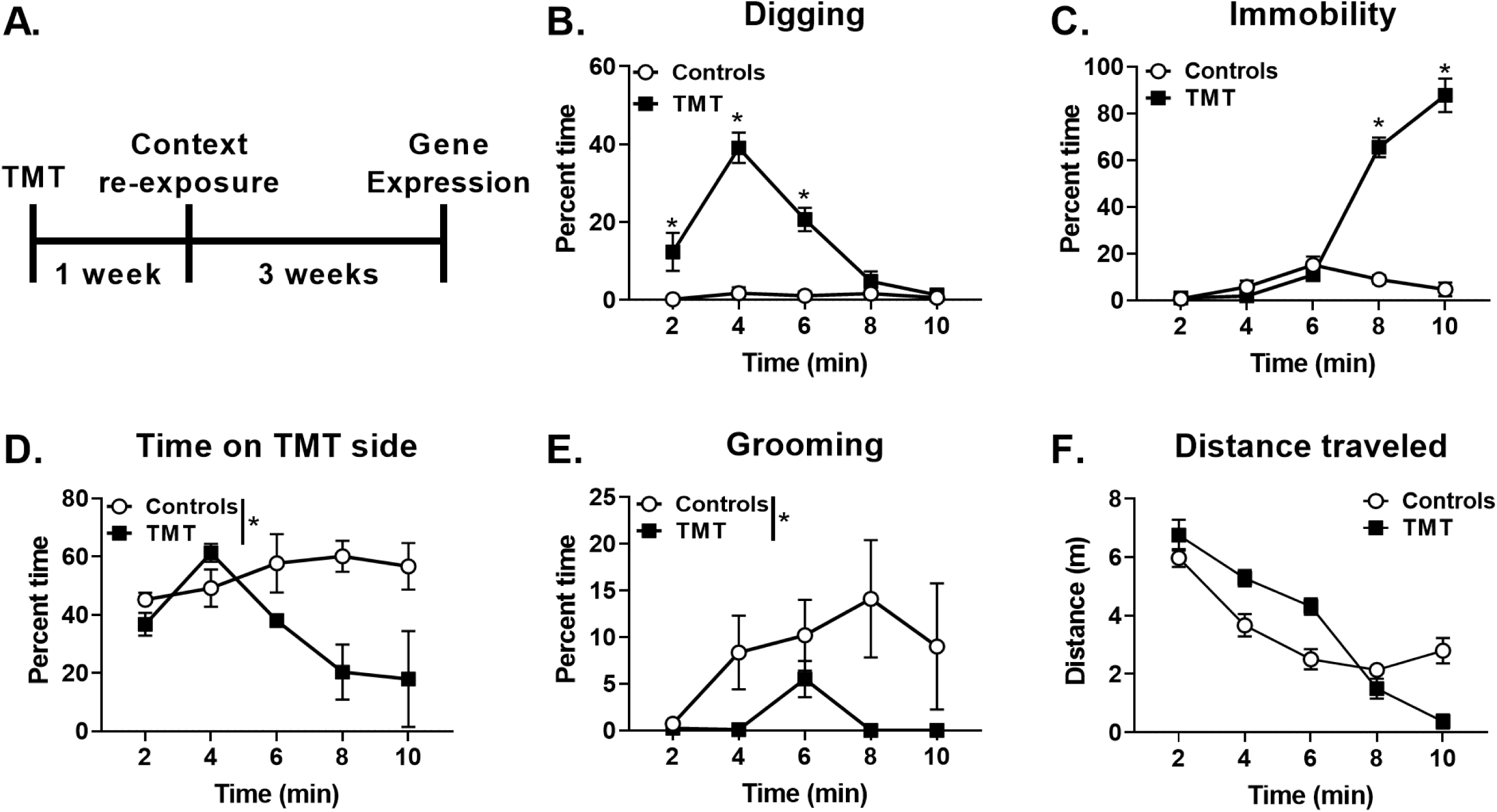
TMT exposure produces digging, freezing, avoidance, and diminished grooming behavior. 3A shows the experimental timeline for Experiment 2. During TMT exposure, the TMT group displayed (B) increased percent time digging, (C) increased percent time immobile, (D) decreased percent time spent on the TMT side, (E) decreased percent time spent grooming, and (F) no change in distance traveled. n=6 Control; n=6 TMT. * p≤0.05 significantly different from Control.

#### Context Re-exposure – enhanced behavioral reactivity to the TMT-paired context

Figure 4 shows the behavioral profile during re-exposure to the previously paired TMT context one week following TMT exposure. No group differences were observed for time spent digging during context re-exposure (Fig. 4A). However, the TMT group displayed decreased immobility (t(14)=2.2, p=0.04, Fig. 4B), increased time spent on the TMT side of the chamber (t(14)=3.5, p=0.004, Fig. 4C), and less time grooming (t(14)=3.3, p=0.006, 4D) than controls. TMT exposure did not affect total distance traveled during context re-exposure (Control: 9.0 ± 0.7 m; TMT: 9.4 ± 0.6 m, not shown). Lastly, a trend for increased fecal boli production was observed in the TMT group compared to controls (Control: 0.6 ± 0.3; TMT: 2.6 ± 0.94; t(14)=2.0, p=0.06). Together, these data show a behavioral reactivity to the TMT-paired context.

**Figure 4.**
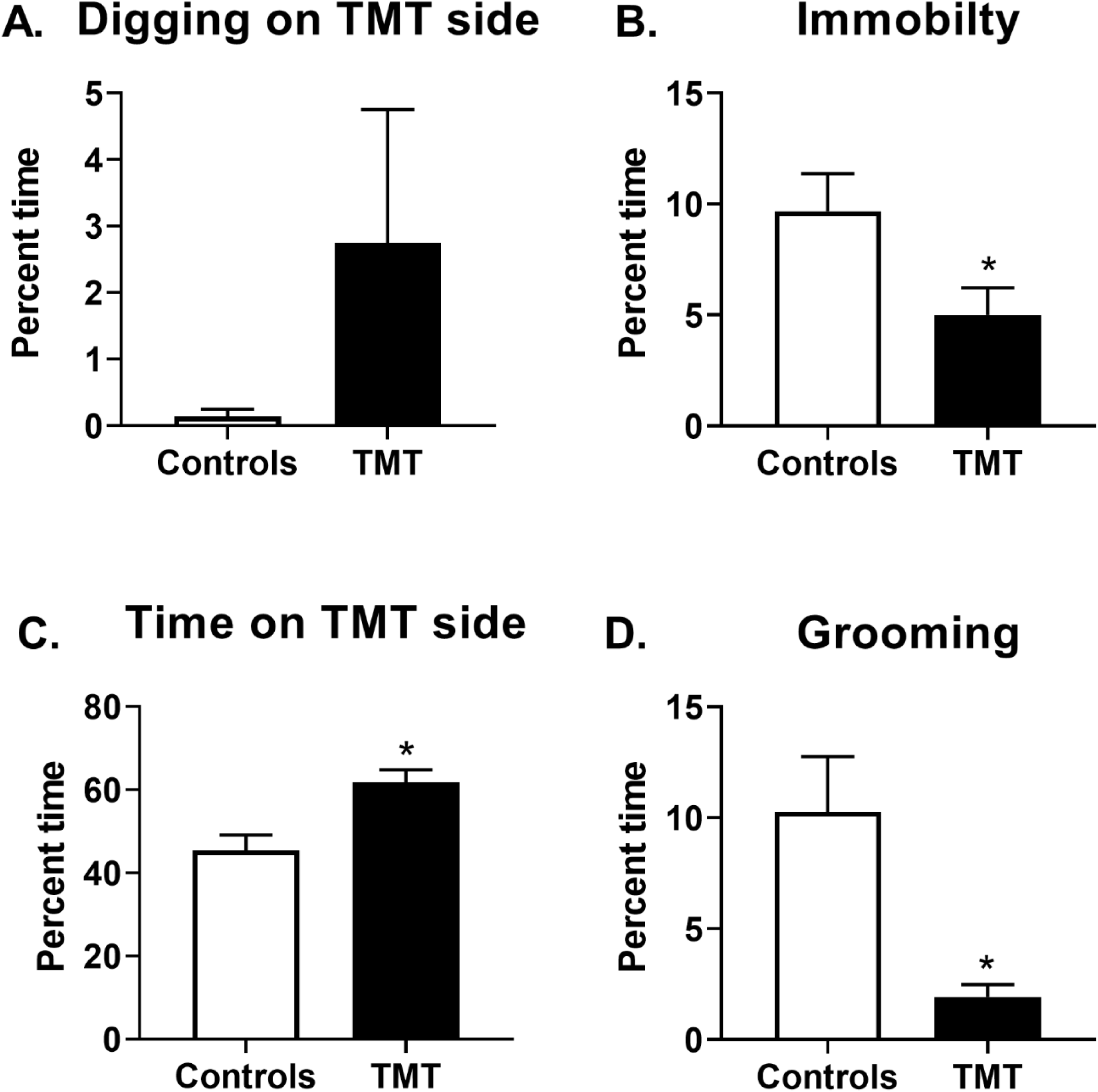
The TMT group displayed context re-exposure behavioral reactivity. One week after TMT exposure, (A) no significant difference in digging behavior between the TMT and control group was observed. However, (B) the TMT groups showed less time spent immobile, (C) more time spent on the TMT side of the chamber, and (D) less time grooming relative to controls. n=8 Control; n=8 TMT. * p≤0.05 significantly different from Control.

#### Gene expression changes four weeks after TMT exposure

Figure 5 shows all significant gene expression changes four weeks after TMT exposure. Figure 5A illustrates upregulation of the stress-related genes, *FKBP5* (t(13)=2.2, p=0.04) and *NR3C2* (t(13)=2.8, p=0.02), but not *NR3C1* in the dorsal hippocampus compared to controls. Figure 5B shows the effects of TMT exposure on the presynaptic, inhibitory receptor gene targets *GRM2, GRM7*, and *CNR1* in the infralimbic cortex, ventral hippocampus and nucleus accumbens. *GRM2* was decreased in the infralimbic cortex (t(13)=3.924, p=0.002), but *GRM7* and *CNR1* were not affected. In the ventral hippocampus, *GRM7* (t(13)=2.502, p=0.02) and *CNR1* (t(12)=2.930, p=0.01) were both downregulated compared to controls. Similarly, the nucleus accumbens showed decreased *GRM7* (t(14)=2.5, p=0.03) gene expression. Figure 5C shows changes in expression of the synaptic glutamate recycling gene *SLC1A3* with an elevation in the prelimbic cortex (t(12)=3.0, p=0.01), and a decrease in the nucleus accumbens (t(13)=3.0, p=0.01). A trend for elevated *SLC1A3* expression in the dorsal hippocampus was observed (p=0.06). Figure 5D shows changes in excitatory post-synaptic density and function (*GRM5, DLG4, Homer1*, and *SHANK3*) gene expression in prelimbic cortex, infralimbic cortex, ventral hippocampus and dorsal hippocampus. In the prelimbic cortex, a nearly 13-fold increase in *GRM5* (t(12)=41.6, p<0.0001) expression was observed in the TMT group compared to controls. This was accompanied by an increase in prelimbic *DLG4* (t(13)=2.7, p=0.017), and a trend for an increase in prelimbic *Homer1* (t(13)=2.0, p=0.07). The infralimbic cortex did not show changes in *GRM5* expression, but did show increased *Homer1* (t(14)=2.2, p=0.05) in the TMT group. In contrast to the prelimbic cortex, the ventral hippocampus showed decreased *GRM5* expression (t(13)=2.4, p=0.03), as well as decreased *SHANK3* (t(13)=4.1, p=0.001) and *Homer1* (t(14)=2.6 p=0.02) in the TMT group compared to controls. Similar to the ventral hippocampus, the dorsal hippocampus showed decreased *SHANK3* expression (t(13)=4.6, p=0.0005), but *GRM5* was not significantly affected.

**Figure 5.**
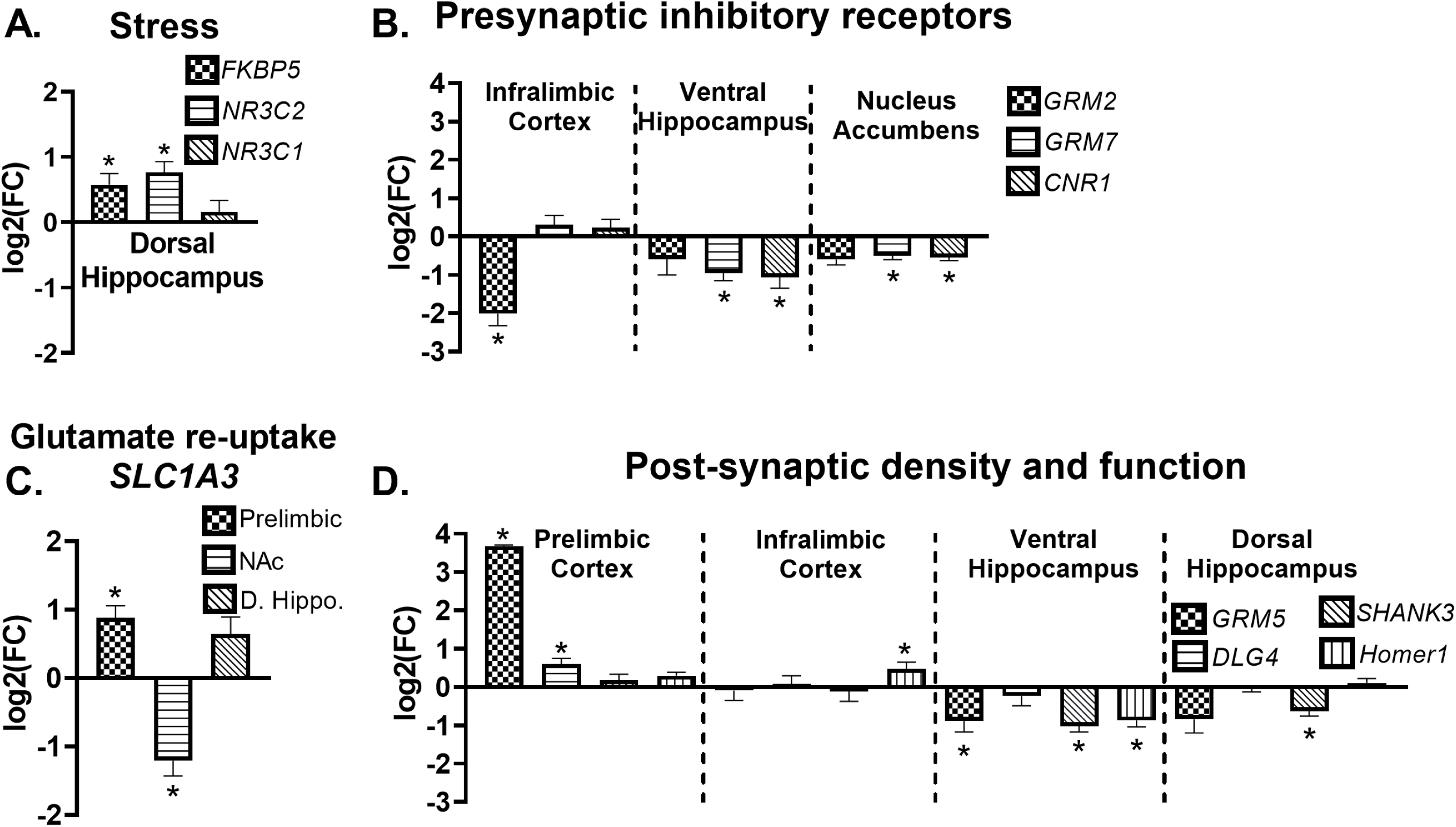
Gene expression changes four weeks after TMT exposure. Four weeks after TMT, (A) *FBKP5* (n=7 Control; n=8 TMT) and *NR3C2* (n=7 Control; n=8 TMT) gene expression were upregulated in the dorsal hippocampus. (B) *SLC1A3* was upregulated in the prelimbic cortex (n=6 Control; n=8 TMT), and decreased in the nucleus accumbens (n=7 Control; n=8 TMT). (C) *GRM2* was downregulated in the infralimbic cortex (n=7 Control; n=8 TMT), *GRM7* (n=8 Control; n=7 TMT) and *CNR1* (n=7 Control; n=7 TMT) were decreased in the ventral hippocampus, and *GRM7* downregulated in the nucleus accumbens (n=8 Control; n=8 TMT). (D) *GRM5* (n=7 Control; n=7 TMT) and *DLG-4* (n=7 Control; n=8 TMT) were upregulated in the prelimbic cortex, and *Homer1* increased in the infralimbic cortex n=7 Control; n=8 TMT). *GRM5* (n=8 Control; n=7 TMT), *Homer1* (n=8 Control; n=7 TMT) and *SHANK3* (n=8 Control; n=7 TMT) were decreased in the ventral hippocampus, and dorsal hippocampus also showed (D) decreased *SHANK3* expression (n=7 Control; n=8 TMT). * p≤0.05 significantly different from Control.

#### Percentage of differentially expressed genes at each time point examined (6 hrs, 2 d, 4 wks)

Figure 6 displays the number of differentially expressed genes as a percentage of all gene expression experiments (gene x ROI) conducted at each time point. 4.7% (2 out of 43) of gene expression examined were differentially expressed six hours post-TMT. 7.0% (3 out of 43) of gene expression examined were differentially expressed 2 days post-TMT. In contrast, four weeks after TMT showed that 20.9% (9 out of 43) of gene expression experiments conducted yielded differential expression. Only those genes and brain regions that were assessed at all three time points were included in these analyses, resulting in a total of 43 experiments at each time point.

**Figure 6.**
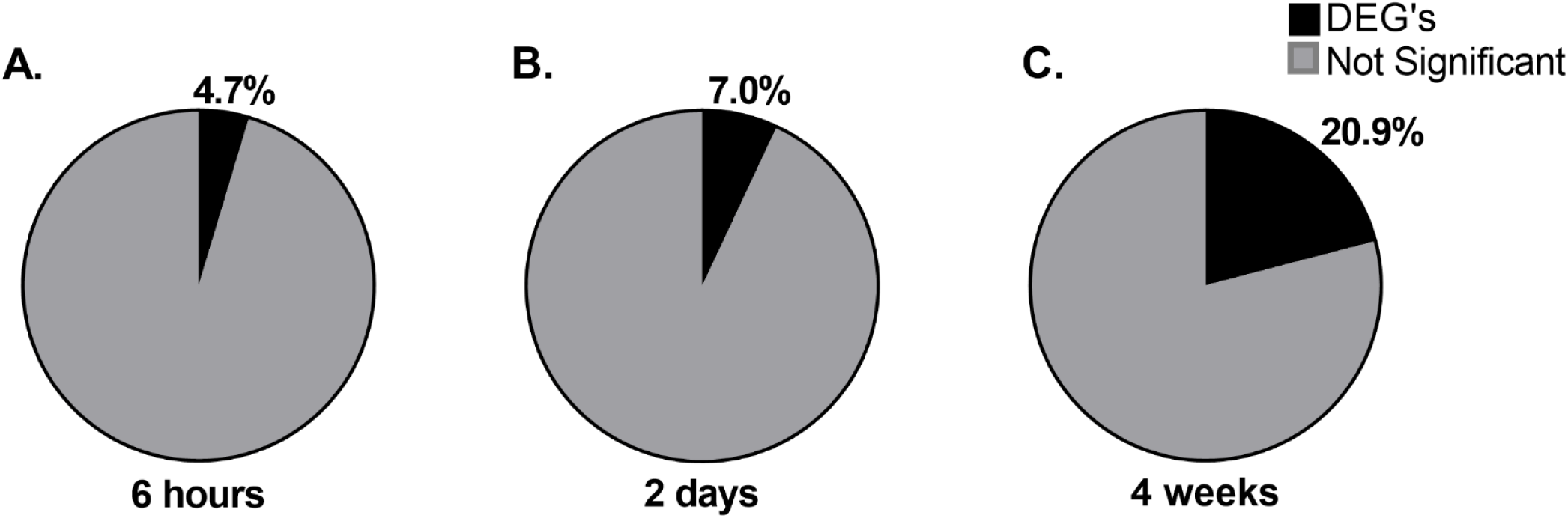
Percentage of differentially expressed genes across the time points. The percentage of differentially expressed genes that were examined at each time point was calculated as 4.7% 6 hours after TMT, 7.0% 2 days after TMT, and 20.9% 4 weeks after TMT. N=43 experiments per time point.

## Discussion

These data demonstrate that exposure to the predator odor TMT induces both early (6 hours and 2 days) and late (4 weeks) gene expression changes related to stress and excitatory synaptic function. Among other targets, TMT exposure affected brain gene expression of *FKPB5, GRM5*, and *CNR1*, which have all been implicated in PTSD^16,20,22^. Furthermore, most differentially expressed genes were observed at late (20.9%) compared to early time points (4.7-7.0%; Figure 6).

First, we established that TMT exposure produced a behaviorally-defined stress response. In both Experiments 1 and 2, rats engaged in immobility and avoidance (decreased time spent on the TMT side) behavior. Note that immobility was operationally defined as lack of movement for more than 2 seconds as assessed using ANY-maze software. Therefore, immobility captures both inactivity and freezing, which is characteristic of a fear response in rodents^39^. Importantly, during the last 2 mins of TMT exposure, the TMT group was immobile for approximately 90% of the time, compared to 9% for controls. In Experiment 2, the addition of bedding to the chamber enabled quantification of an additional stress-reactive behavior - digging. Rats engaged in digging behavior during the first 6 mins of exposure, which ultimately resulted in a pile of bedding under the TMT source. Digging behavior appears similar to defensive burying behavior, which rodents display in response to predator odors^40^. However, because hanging the predator odor from a basket makes burying impossible, digging may reflect a failed attempt at defensive burying behavior. By min 8, the TMT group began to engage in freezing and avoidance behavior similar to Experiment 1. Results from Experiment 1, which used a longer, 20 min TMT exposure, support the observation that once rats begin engaging in freezing and avoidance behaviors, they remain that way until the termination of the experiment. Finally, control rats spent significantly more time grooming than the TMT group, presumably because the TMT group was engaging in digging and freezing behaviors. Increased digging, immobility, avoidance, and decreased grooming during the TMT exposure are consistent with previous findings using TMT stress^26,40,30,41^. Together, these data reflect a shift from an early, active approach behavior (digging) to a passive immobility/freezing and avoidance behavior.

In Experiment 2 one week after TMT, rats were re-exposed to the context in which they were exposed to TMT in order to assess for a lasting behavioral response to the TMT-paired context. Rats in the TMT group showed an altered behavioral response to the TMT-paired context relative to controls. This was displayed by behaviors indicative of exploration and increased activity, such as decreased immobility, increased time spent in the TMT side of the chamber, and decreased time spent grooming. In contrast to the immobility observed during TMT exposure (previously discussed), immobility during the context re-exposure likely reflects inactivity, and not freezing, especially given that the TMT group spent only 5% of the total time immobile (controls: 10%). Therefore, a possible interpretation of decreased immobility (i.e., increased mobility) and grooming in the TMT group during context re-exposure is that control rats showed greater habituation to the context (more inactivity, more grooming), whereas rats that previously experienced the context paired with TMT engaged in more exploratory behavior (less inactivity, less grooming), which may reflect stress-reactivity or hypervigilance to the TMT-associated context. Surprisingly, the TMT group spent slightly more time on the TMT side of the chamber than controls. This could reflect an exploratory behavior of the basket that previously contained TMT or greater exploration of the TMT side in general as this was the side that was previously avoided and thus less explored on the initial exposure. Together, these data reflect a lasting behavioral reactivity to the TMT-paired context in rats previously exposed to TMT.

TMT exposure resulted in both immediate (6 h and 2 days) and late changes (4 weeks) in stress-related genes. *FKBP5* gene expression was increased in the hypothalamus and dorsal hippocampus 6 hours following TMT, but returned to control values 2 days after TMT. Additionally, *CRF* mRNA trended upwards (p=0.06) 2 days after TMT exposure in the hypothalamus. Together, these data suggest engagement of the HPA-axis, and peripheral glucocorticoid release in response to TMT exposure^42^. Interestingly, *FKBP5* was also upregulated 4 weeks after TMT in the dorsal hippocampus, which showed increased *NR3C2* (encoding MR), but not *NR3C1* (encoding GR), expression as well. FKBP5 forms a complex with the GR and MR, preventing their translocation into the nucleus, and thus reduces GR- and MR-mediated transcriptional regulation of HPA-axis negative feedback^13^. Therefore, binding of FKBP5 to GR and MR might counteract these receptors’ effects on HPA-axis negative feedback. Interestingly, PTSD is associated with altered *FKBP5* transcription and functionally relevant changes in FKBP5-GR complex^16,43^. These data suggest that the TMT exposure model of stress could be used to investigate the role of FKBP5 in both early and late effects of traumatic stress.

Stress-induced increases in glucocorticoid levels affect multiple aspects of glutamate/excitatory brain neurotransmission, indicating a functional relationship between stress and excitatory synaptic transmission^24^. Therefore, finding of changes in the stress-related genes were followed up by examining genes known to modulate glutamate neurotransmission. At early time points, we identified differential expression of *GRM3* across three brain regions. At long-term time points, we showed that genes known to mediate presynaptic neurotransmitter release (i.e., *GRM2, GRM7, CNR1*), synaptic glutamate recycling (i.e., *SLC1A3*), and post-synaptic excitatory signaling (i.e., *GRM5, DLG4, Homer1, SHANK3*) were differentially expressed following TMT exposure.

Two days after TMT exposure, *GRM3* (encoding mGluR3) was decreased in the prelimbic cortex and dorsal hippocampus, but increased in the nucleus accumbens, with no significant changes in *GRM2, GRM5* and *GRM7* expression in any brain region examined at early time points. *GRM3* is part of Group II metabotropic glutamate receptors (mGluR2 and 3), which are coupled to Gi/Go to negatively regulate adenylyl cyclase activity^44^. These receptors are considered largely presynaptic, acting as inhibitory receptors to diminish neurotransmitter release^44^. However, mGluR3 is also expressed at the post-synaptic membrane and on astrocyte projections at the tripartite synapse^44^, and displays a post-synaptic site of function in the prelimbic cortex^45^. Group II mGluRs play an important role in stress, with several studies demonstrating a functional role of these receptors in stress-induced anhedonia/depressive- and anxiety-like behavioral phenotypes^44, 45, 46,47,48^. Negative allosteric modulation of mGluR3 has antidepressant-like effects in the forced swim and marble burying tests^49^. Interestingly, mGluR3-mediated long-term depression (LTD) on excitatory prelimbic neurons was abolished following restraint stress^45^, which is consistent with our findings that TMT stress downregulates *GRM3* expression in the prelimbic cortex. Therefore, the observed effects of predator odor stress on *GRM3* gene expression may reflect stress-induced changes to synaptic plasticity that could in part underlie the observed changes in subsequent context re-exposure behavioral reactivity and gene expression changes related to the excitatory synapse^50^.

Four weeks after TMT exposure, several changes in genes encoding Gi/o-coupled presynaptic glutamate receptors were observed. *GRM2* expression was decreased in the infralimbic cortex (Il). As previously stated, mGluR2 receptors are presynaptic, inhibitory receptors^44^ that have been implicated in stress susceptibility^51^. Additionally, *GRM7* (encoding mGluR7) and *CNR1* (encoding CB1) were decreased in the ventral hippocampus, and *GRM7* decreased in the nucleus accumbens. mGluR7 is part of Group III mGluRs, which are Gi/o coupled GPCRs, reducing cyclic AMP formation similar to that of Group II mGluRs, but show different affinity for glutamate compared to Group II receptors^52^. mGluR7 displays low affinity for glutamate, and activation reduces glutamate release under conditions of high extracellular glutamate concentrations^52^. Additionally, the CB1 receptor is a presynaptic receptor that when activated by endocannabinoids inhibits neurotransmitter release, and has been implicated in stress and PTSD^22,53^. Together, these results demonstrate that downregulation of presynaptic, inhibitory receptor gene expression in infralimbic cortex, ventral hippocampus, and nucleus accumbens are persistent consequences of TMT stress, suggesting possible late effects on extracellular glutamate concentrations.

To follow-up on these results, we examined expression of *SLC1A3* (encodes the EAAT-1 receptor) as a more direct indicator of changes in extracellular glutamate concentrations. *SLC1A3* gene expression was increased in the prelimbic cortex and decreased in the nucleus accumbens. EAAT-1 is expressed on astrocyte projections at the tripartite synapse, and serves to recycle extracellular glutamate from the synapse^54^. Interestingly, PTSD is associated with high extracellular glutamate, and changes in other neurotransmitter concentrations^18^. These data provide further support for the hypothesis that TMT exposure induced late changes in excitatory molecular composition related to synaptic glutamate levels.

Next, we investigated whether these changes in presynaptic genes and extracellular glutamate markers were accompanied by changes in the post-synaptic glutamate receptor, *GRM5* (encoding mGluR5). *GRM5* was differentially expressed in both the prelimbic cortex (increased) and ventral hippocampus (decreased). mGluR5 receptors are part of Group I mGluRs that are Gq/s coupled, increasing cAMP activity on the postsynaptic cell, and affecting NMDAR-mediated excitability^55^. Additionally, preclinical studies have demonstrated a functional role for mGluR5 signaling in stress-related behaviors^31,56,57^. Interestingly, in a clinical study in individuals with PTSD, cortical mGluR5 availability was increased relative to healthy controls, and positively correlated with avoidance symptom severity^20^. Therefore, the nearly 13-fold increase in *GRM5* expression in the prelimbic cortex observed in Experiment 2 suggests that increased stress-related cortical mGluR5 might be transcriptionally regulated, and may be a conserved, long-term adaptation that persists following a severe stressor. In contrast to the prelimbic cortex, the ventral hippocampus showed decreased expression of *GRM5*, suggesting the possibility that *GRM5* plays a distinct stress-related role depending on the brain region. Group I mGluR agonism induces robust electrophysiological readouts particularly in the ventral hippocampus, which may play a role in fear memory or extinction mechanisms^58,59^.

To follow-up on the changes in *GRM5*, we examined gene expression of intracellular post-synaptic targets that complex with glutamate membrane receptors (*DLG4, SHANK3*, and *Homer1*) in brain regions where *GRM5* was changed (prelimbic cortex and ventral hippocampus)^34^. Further, we examined the dorsal hippocampus and infralimbic cortex to determine if changes were specific to the ventral hippocampus and prelimbic cortex. Interestingly, at least one of these post-synaptic density genes (*DLG4, SHANK3, Homer1*) were changed in regions where *GRM5* was also affected. Further, these gene expression changes were in the same directionality of expression as *GRM5*. Specifically, *DLG4* was increased in the prelimbic cortex, *Homer1* increased in the infralimbic cortex, *SHANK3* and *Homer1* decreased in the ventral hippocampus, and *SHANK3* decreased in the dorsal hippocampus. Together, these results suggest that TMT exposure produced late changes in gene expression related to excitatory post-synaptic signaling.

As previously mentioned, predator odor stress models have been used to behaviorally - define stress-”susceptible” and “resilient” groups in an effort to improve the face validity of predator odor models^25,29,32^. However, the goal of the present experiments were to provide a comprehensive and dynamic assessment of glutamate- and stress-related genes throughout the brain following a single exposure to the predator odor TMT. To this end, the sample sizes used were not sufficient to define sub-populations, but it will be important for future work to define how the genes identified here play a role in resilience and vulnerability to develop maladaptive behavioral changes. Another consideration is that these studies were conducted in male rats only. Sex differences in response to predator odor are documented^60^ and women suffer from PTSD at three times the rate of men^3,16^. Therefore, future work should examine sex differences in response to TMT stress.

While conclusions about function cannot be determined based on gene expression data alone, the magnitude of gene expression changes related to a similar function (i.e., excitatory/glutamate signaling and the stress response) suggests the strong possibility that some adaptation to stress mechanisms and excitatory signaling occurred following TMT stress. These data show how TMT stress affects early and late glutamate- and stress-related gene expression throughout the brain relevant to targets identified in the clinical literature. More changes were observed at late as compared to early time points, providing temporal information about the nature of TMT-induced molecular adaptations. These data build upon our understanding of the molecular changes following a predator odor stressor, which could inform our understanding of traumatic stress disorders.

## Acknowledgements

This work was supported in part by the National Institute of Health AA026537 (JB) and by the Bowles Center for Alcohol Studies. RET was supported by NS007431. The authors thank Abigail Garcia-Baza for her help with behavioral analyses.

## Data Availability Statement

Data available on request from the authors.

