## Supplemental Information for "Exposure to the predator odor TMT induces early and late differential gene expression related to stress and excitatory synaptic function throughout the brain in male rats"

**Supplementary Table 1 – Behavioral analyses in seconds**

| TMT Exposure (10 min, s) | | | | |
| --- | --- | --- | --- | --- |
|  | Digging | Immobility | Time on TMT side | Grooming |
| Controls | 6.3 ± 3.5 | 42.0 ± 8.2 | 322.4 ± 15.6 | 50.8 ± 8.7 |
| TMT | 93.8 ± 15.5* | 200.3 ± 7.3* | 209.0 ± 31.6* | 7.0 ± 0.3* |
| Context Re-exposure (5 min, s) | | | | |
|  | Digging | Immobility | Time on TMT side | Grooming |
| Controls | 1,4 ± 1.0 | 29.0 ± 5.1 | 136.0 ± 11.3 | 30.8 ± 7.5 |
| TMT | 8.3 ± 6.0 | 15.0 ± 3.7* | 185.5 ± 8.8* | 5.8 ± 1.7* |

S1. Supplementary Table 1 – The total time (s) spent engaging in each behavior during TMT exposure and context re-exposure for Experiment 2. * p<0.05 vs. Control (p≤0.05).

**Supplementary Table 2 – Gene expression 6 hours after TMT**
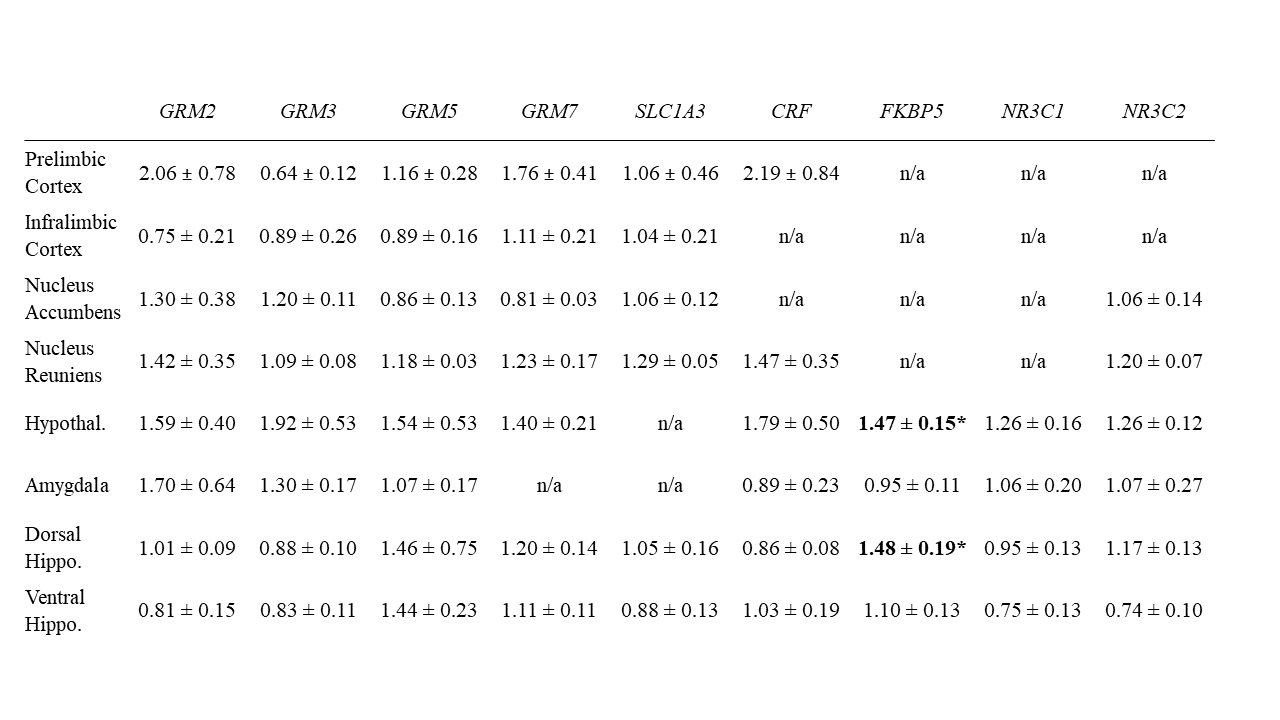
S2. Supplementary Table 2 – A complete list of gene expression data as fold changes of TMT group compared to control (average=1) 6 hours after TMT exposure. * p≤0.05 vs. Control

**Supplementary Table 3 – Gene expression 2 days after TMT**

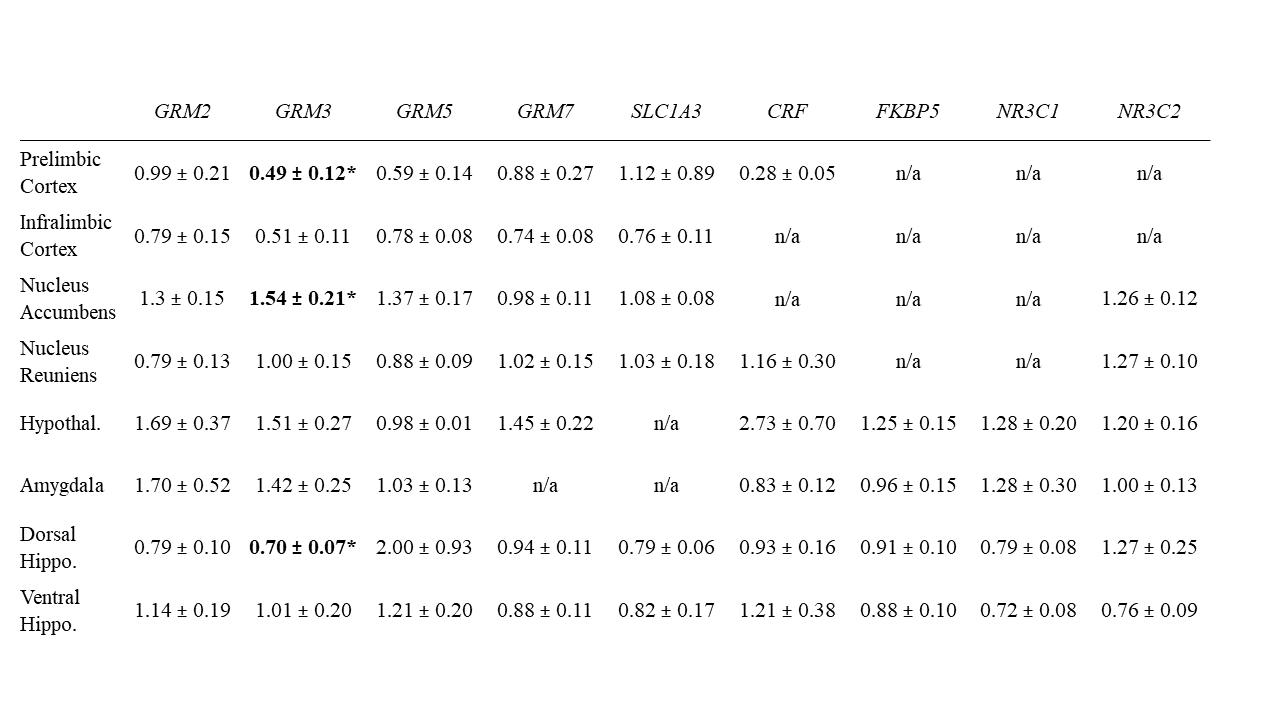
S3. Supplementary Table 3 - A complete list of gene expression data as fold changes of TMT group compared to control (average=1) 2 days after TMT exposure. Data from S2 and S3 are from Experiment 1. * p≤0.05 vs. Control.

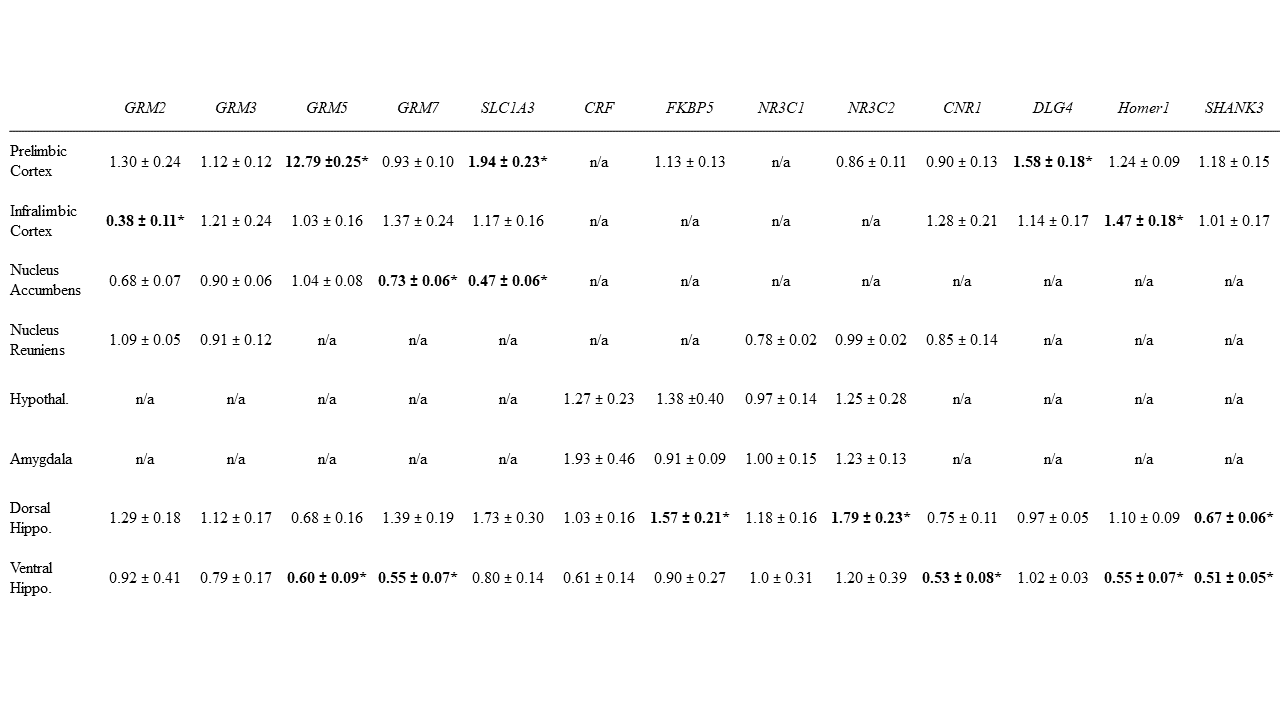
**Supplementary Table 4 – Gene expression 4 weeks after TM**T

Supplementary Table 4 - A complete list of gene expression data as fold changes of TMT group compared to control (average=1) 4 weeks after TMT exposure. Data from S4 are from Experiment 2. * p≤0.05 vs. Control.
